# Dietary *S. maltophilia* promotes fat storage by enhancing lipogenesis and ER-LD contacts in *C. elegans*

**DOI:** 10.1101/2020.04.29.067793

**Authors:** Kang Xie, Yangli Liu, Xixia Li, Hong Zhang, Shuyan Zhang, Ho Yi Mak, Pingsheng Liu

**Affiliations:** National Laboratory of Biomacromolecules, CAS Center for Excellence in Biomacromolecules, Institute of Biophysics, Chinese Academy of Sciences, Beijing 100101, China; University of Chinese Academy of Sciences, Beijing 100049, China; Division of Life Science, The Hong Kong University of Science and Technology, Hong Kong SAR, China

**Keywords:** *Stenotrophomonas maltophilia*, obesity, DHS-3, MDT-28, lipid droplet

## Abstract

Dietary and symbiotic bacteria can exert powerful influence on metazoan lipid metabolism. Here, we demonstrate that feeding *Caenorhabditis elegans (C. elegans)* with the opportunistic pathogenic bacteria *Stenotrophomonas maltophilia (S. maltophilia)* retards growth and promotes excessive fat storage. Gene expression analysis reveals that dietary *S. maltophilia* induces a lipogenic transcriptional response that includes the SREBP ortholog SBP-1, and fatty acid desaturases FAT-6 and FAT-7. Live imaging and ultrastructural analysis suggest that excess fat is stored in greatly expanded lipid droplets (LDs), as a result of enhanced endoplasmic reticulum-LD interaction. We also report that loss of function mutations in *cyp-35B1* or *dpy-9* in *C. elegans* confers resistance to *S. maltophilia.* Our work delineates a new model for understanding microbial regulation of metazoan physiology.

## Introduction

In the wild, the nematode *C. elegans* feeds on a variety of soil bacteria including *Pseudomonas medocina, Bacillus megaterium, Comomonas sp.* (Avery & Shtonda, 2003, Duveau & Felix, 2012, Felix & Duveau, 2012, Montalvo-Katz, Huang et al., 2013, Watson, MacNeil et al., 2014b, Zhang, Holdorf et al., 2017). In contrast, the *E. coli* OP50 is commonly used as the standard laboratory diet (Brenner, 1974, Sulston & Brenner, 1974). Therefore, the full range of physiological response in *C. elegans* toward dietary bacteria remains obscure. Nevertheless, increasing evidence suggests that *C. elegans* is capable of integrating olfactory, mechanical and nutritional cues to identify its preferred bacterial diet. Such dietary choices can subsequently exert significant impact on *C. elegans* lifespan and metabolism (Brooks, Liang et al., 2009, Garcia-Gonzalez, Ritter et al., 2017, Gusarov, Gautier et al., 2013, Han, Sivaramakrishnan et al., 2017, Lin & Wang, 2017, MacNeil, Watson et al., 2013, Qi & Han, 2018, Scott, Quintaneiro et al., 2017, Virk, Correia et al., 2012, Watson, MacNeil et al., 2014a).

Lipid droplets (LDs) are conserved organelles for cellular fat storage. Over the last decade, our laboratory has focused on using mass spectrometry to determine the proteome of LDs that were biochemically purified from a wide range of uni- and multi-cellular organisms, including *C. elegans.* Accordingly, we identified two major LD resident proteins in *C. elegans,* DHS-3 and MDT-28 (Liu, Xu et al., 2018, Na, Zhang et al., 2015, Zhang, Na et al., 2012). When expressed as fluorescent fusion proteins in transgenic worms, DHS-3 and MDT-28 serve as faithful marker for monitoring LD morphology in live animals. Importantly, we found a strong correlation between LD size and number with organismal fat storage.

Here we apply our suite of fluorescence markers for the ER and LDs, to study the effect of an environmental bacterium *S. maltophilia* on fat storage in *C. elegans*. Interestingly, clinical isolates of *S. maltophilia* have been regarded as pathogenic in immunocompromised humans. We found that *C. elegans* grew slower and accumulated significantly more fat when fed *S. maltophilia,* instead of the standard laboratory diet*, E. coli* OP50. By genetic analysis, we uncovered a transcriptional regulatory network that promoted lipogenesis in response to *S. maltophilia.* Coupled with the remodeling of ER-LD interaction, massive LD expansion ensued. Accordingly, attenuation of ER-LD interaction partially suppressed the effect of dietary *S. maltophilia* on fat storage. Our results help establish a new *C. elegans*-microbe experimental paradigm for the study of dietary factors that modulate lipid metabolism.

## Results

### Lipid Droplet Size in *C. elegans* Is Dependent on Dietary Microbe

Two bacterial species were previously shown to reduce *C. elegans* lipid accumulation (Ding, Smulan et al., 2015, Kishino, Takeuchi et al., 2013). To recapitulate these results, we fed *Lactobacillus plantarum* and *Pseudomonas aeruginosa* (strain UCBPP-PA14, PA14) to transgenic worms that expressed the intestinal LD marker, DHS-3::GFP. We found that *Lactobacillus plantarum* and *Pseudomonas aeruginosa* reduced the average LD diameter by 26.7% and 83.3%, respectively (Figures 1A-E). We were therefore encouraged to use our DHS-3::GFP reporter strain for further exploration of the effect of bacterial diets on *C. elegans* fat storage (Brooks et al., 2009).

**Figure 1.**
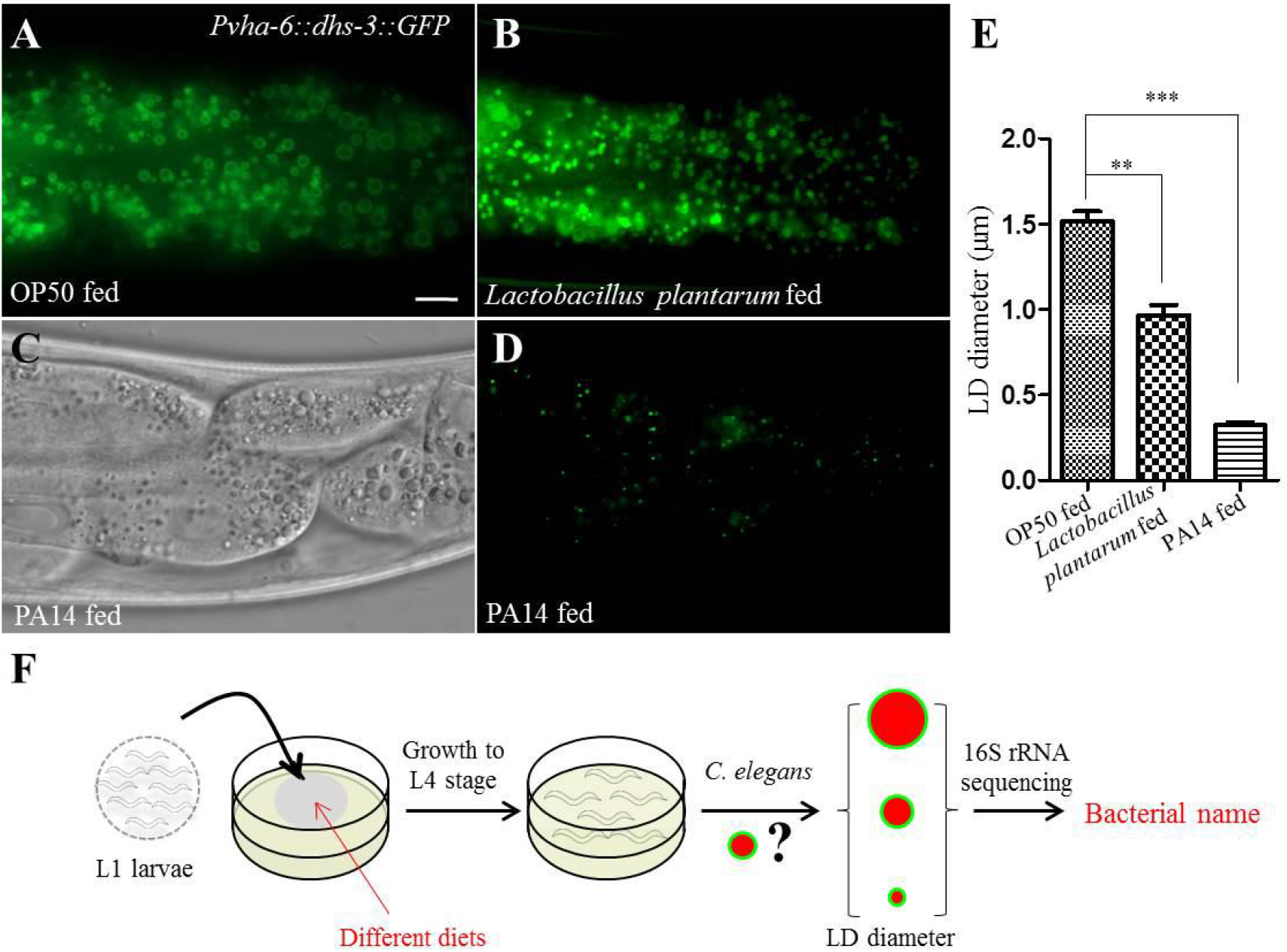
*Lactobacillus plantarum* and PA14 reduce the size of LDs in *C. elegans.* **(A**) Fluorescence micrographs of LDs in the intestine of OP50-fed worms. Scale Bar, 5μm. **(B)** Fluorescence micrographs of *Pvha-6::dhs-3::GFP* in *Lactobacillus plantarum-fed* worms. **(C)** Differential interference contrast (DIC) images of LDs in PA14-fed worms. **(D)** Fluorescence micrographs of *Pvha-6::dhs-3::GFP* in PA14-fed worms. **(E)** Quantification of the LD diameter (A, B, D). Data represent mean ± SEM (n=5 for each independent experiment, \*\**P*<0.01, \*\*\**P*<0.001, one-way ANOVA). **(F)** Schematic representation of the screening method to identify bacteria that affect host LDs.

We performed a screen to identify bacteria that modulate *C. elegans* lipid metabolism. To capture environmental bacteria, Nematode Growth Medium (NGM) plates were left open in the laboratory. Bacteria that yielded individual colonies were isolated, cultured in liquid medium, and subsequently re-introduced to NGM plates to form single-species bacterial lawns. Synchronized L1 stage transgenic reporter worms that express DHS-3::GFP were allowed to develop with the environmental bacteria as food. We subsequently quantified the LD size in larval L4 stage worms. Bacteria that significantly altered LD size of worms were taken through multiple cloning steps to obtain a pure clone. The cultures were then fed again to worms to verify the effect and the bacteria were identified by 16S rRNA sequencing (Fig. 1F). Worms fed these positive clones invariably showed growth retardation or arrest (Table 1). We then focused on *Stenotrophomonas maltophilia* because it was the only bacterial species that significantly increased the LD size of worms.

**Table 1.**
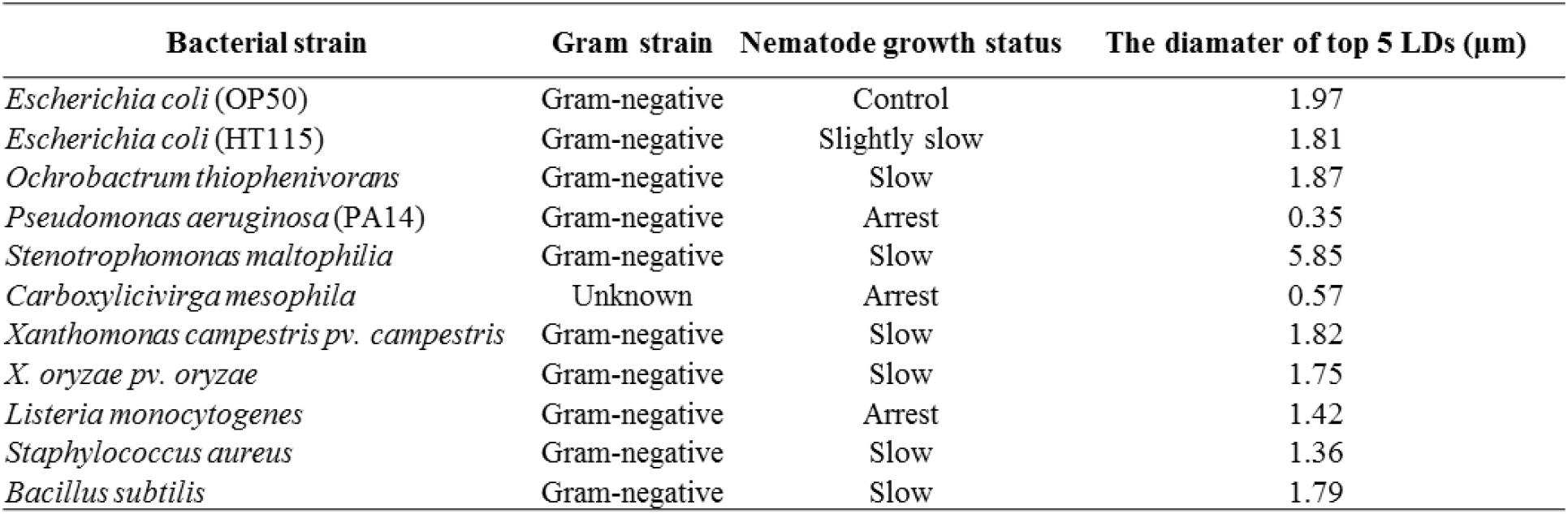
The effects of different bacterial diets on *C. elegans* growth and LD size.

### *Stenotrophomonas maltophilia* promotes LD expansion in *C. elegans*

The bacterium *Stenotrophomonas maltophilia* induced a striking increase in LD diameter after 2 days of feeding (Table 1). To confirm the effect, we repeated the experiment using *S. maltophilia* strains isolated in other laboratories (Liu, Tian et al., 2017, Wang, Li et al., 2014) (Table 2). Similar to our original isolate, all strains tested reproducibly promoted LD expansion in worms (Fig. 2A). Our results suggest that *S. maltophilia* harbors a species-specific factor that potently modulates fat storage in *C. elegans*.

**Figure 2.**
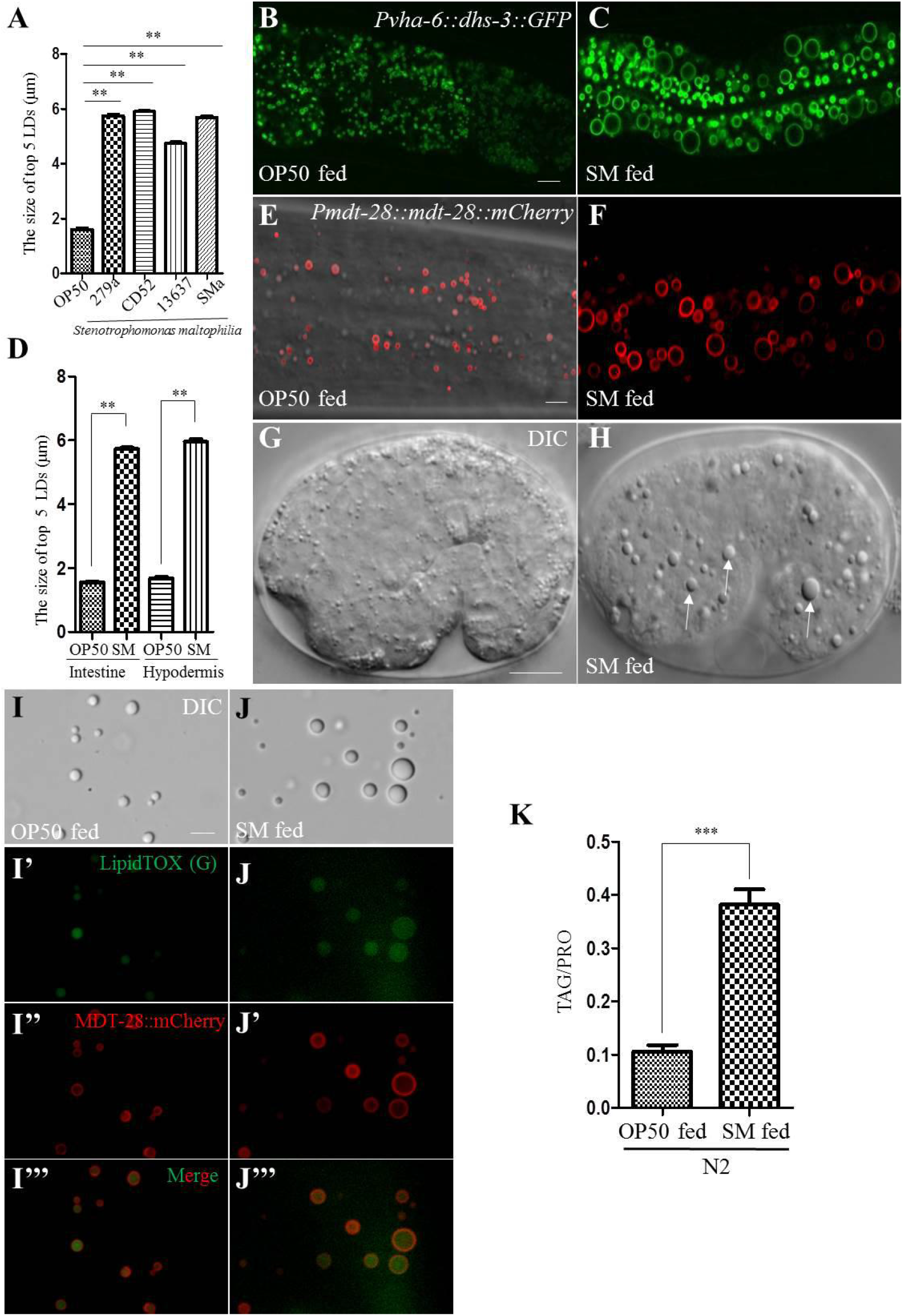
*Stenotrophomonas maltophilia* induces supersized LDs in *C. elegans.* **(A)** Quantification of LD diameter in N2 worms fed different *S. maltophilia* strains. Data represent mean ± SEM (n=5 for each independent experiment, \*\**P*<0.01, one-way ANOVA). **(B)** Fluorescence micrographs of LDs in the intestine of OP50-fed worms. Scale Bar, 5 μm. **(C)** Fluorescence micrographs of *Pvha-6::dhs-3::GFP* in *S. maltophilia-fed* worms. **(D)** Quantification of LD diameter (B, C, E, and F). Data represent mean ± SEM (n=5 for each independent experiment, \*\**P*<0.01, student *t*-test). **(E)** Fluorescence micrographs of LDs in the hypodermis of OP50-fed worms. Scale Bar, 5 μm. **(F)** Fluorescence micrographs of *Pmdt-28::mdt-28::mCherry* labeled LDs in *S. maltophilia-fed* worms. **(G)** DIC images of LDs in the embryonic stage of OP50-fed worms. Scale Bar, 5 μm. **(H)** DIC images of LDs in the embryonic stage of *S. maltophilia*-fed worms. The white arrows point to the *S. maltophilia-enlarged* LDs. **(I)** DIC images of LDs purified from transgenic animals expressing MDT-28::mCherry and fed OP50. Scale Bar, 5 μm. **(I’-I’’)** Fluorescence micrographs of purified LDs from transgenic animals expressing MDT-28::mCherry and fed OP50. LipidTOX Green stained-LDs (I’) and MDT-28::mCherry labeled LDs (I”). **(I’’’)** As in (I’), with LipidTOX (G) and mCherry signals merged. **(J)** DIC images of LDs purified from transgenic animals expressing MDT-28::mCherry and fed *S. maltophilia.* Scale Bar, 5 μm. **(J’-J’’)** Fluorescence micrographs of LDs purified from transgenic animals expressing MDT-28::mCherry and fed *S. maltophilia.* LipidTOX Green stained-LDs (J’) and MDT-28::mCherry labeled LDs (J’’). **(J’”)** As in (J’), with LipidTOX and mCherry signals merged. **(K)** Comparison of TAG levels between OP50 and *S. maltophilia-fed* worms. *** represents *P* value < 0.001, student *t*-test).

**Table 2.** *Stenotrophomonas maltophilia* strains.

| Strain name | Bacterial source |
| --- | --- |
| *Stenotrophomonas maltophilia* 279a | ATCC |
| *Stenotrophomonas maltophilia* CD52 | Keqin Zhang's lab |
| *Stenotrophomonas maltophilia* 13637 | Wei Qian's lab |
| *Stenotrophomonas maltophilia* (s) | Pingsheng Liu's lab |

Next, we asked if the effect of *S. maltophilia* feeding on LD size was restricted to the *C. elegans* intestine. As indicated by the *vha-6* promoter driven, intestine-specific DHS-3::GFP marker, the LD diameter of worms fed *S. maltophilia* was increased 3-fold over those fed OP50 (Fig. 2B-C). To quantify LD size in the hypodermis, we used MDT-28::mCherry that was expressed under the control of the *mdt-28* promoter (Fig. 2E) (Na et al., 2015). Similar to our observations in the intestine, *S. maltophilia* feeding induced LD expansion in the hypodermis. The maximum LD diameter reached 6 μm (Fig. 2D-F). Finally, we compared LD morphology in embryos from parents that were fed either OP50 or *S. maltophilia* (Fig. 2G-H). The embryos from *S. maltophilia*-fed worms again showed enlarged LDs (Fig. 2H, white arrows). These results were confirmed by examining biochemically purified LDs from worms fed OP50 or *S. maltophilia*. The LDs isolated from *S. maltophilia-*fed larval L4 worms, were enlarged relative to OP50-fed worms (Fig. 2I-?” and 2J-J”’). Our results indicate that dietary *S. maltophilia* caused LD enlargement pleiotropically in worms.

The primary cargoes of LDs are neutral lipids, such as triacylglycerol (TAG). To determine if the LD expansion observed in *S. maltophilia* fed worms reflected an increase in organismal lipid content, we directly measured TAG levels biochemically. Total lipids were extracted from worms fed either OP50 or *S. maltophilia* from L1 to L4 stage. We found that *S. maltophilia* fed worms had 3.7-fold more TAG than those fed OP50 (Fig. 2K). Taken together, our imaging and biochemical analyses indicate that *S. maltophilia* can potently induce fat accumulation in *C. elegans*.

### Induction of C. elegans fat storage by S. maltophilia was not due to its species-specific fatty acid composition

Differences in lipid composition between bacterial food sources could potentially underlie the effect of dietary *S. maltophilia* on host lipids, perhaps due to some lipids being more readily absorbed, processed, and incorporated than others. We examined the fatty acid compositions of *E. coli* OP50 and *S. maltophilia* by gas chromatography-mass spectrometry (GC-MS) and found substantial differences (Fig. S1A and S1B). However, it is not clear from this observation alone if lipid compositional difference accounted for the ability of *S. maltophilia* to promote fat accumulation in worms. Therefore, we conducted a transposon-based forward genetic screen in *S. maltophilia* to identify mutants that failed to induce LD expansion when fed to *C. elegans* (Fig. S1C). Through this approach, we isolated SMa9, a mutant *S. maltophilia* strain that could not increase LD size and number in *C. elegans* (Fig. S1D-E). However, there was no difference in fatty acid composition between the mutant SMa9 and the parental wild type (WT) bacteria (Fig. S1F). Therefore, our results clearly suggest that one or more factors other than fatty acids, were responsible for the ability of *S. maltophilia* to promote fat storage in *C. elegans*.

### *S. maltophilia-induced* fat storage was not part of an innate immune response in *C. elegans*

We examined whether the phenotype of excess fat accumulation was due to a pathological process associated with live *S. maltophilia*. Worms were fed ultraviolet (UV)-killed or high temperature-killed *S. maltophilia* and their LDs were examined (Fig. S2A). The killed bacteria retained the ability to increase LD size in *C. elegans,* compared with OP50 feeding. Next, we measured LD size in worms defective in the p38 MAPK pathway *(sek-1, pmk-1),* which is critical for the innate immune response. We found no change in the LD phenotype in these mutant worms (Fig. S2B and S2C). We concluded that the elevated fat storage in worms upon *S. maltophilia* feeding was independent of the innate immune response against pathogens.

### *Stenotrophomonas maltophilia* modulated lipid metabolism in *C. elegans* by Up-regulating *sbp-1, fat-6; fat-7*

We explored the molecular mechanism of *S. maltophilia*-induced LD enlargement in *C. elegans*. To determine the effects of bacteria on *C. elegans* physiology, we measured developmental rate, fecundity, and hatching rate of worms fed either *E. coli* OP50 or *S. maltophilia.* To assess developmental rate, we synchronized animals at the L1 stage and monitored the developmental age of the population over time (Fig. S3A). Development to the L4 stage was delayed in worms fed *S. maltophilia*, compared with OP50 (Fig. S3A). The number of offspring produced by *S. maltophilia*-fed worms was reduced by 30% compared with the OP50 group (Fig. S3B). There was no significant difference in hatching rate between OP50 and *S. maltophilia*-fed worms (Fig. S3C). The *S. maltophilia* bacteria were strongly preferred by worms over the *E. coli* OP50 (Fig. S3D-F). We measured pharyngeal pumping in worms after 1 h and 3 h of feeding and found no differences (Fig. S3G). Lastly, we investigated the effect of *S. maltophilia* feeding on LD size in *eat-4(ky5)* mutant worms, which exhibited reduced pharyngeal pumping and therefore ate less than wild type worms (Hills, Brockie et al., 2004). There were no differences in LD size between wild type and *eat-4(ky5)* mutant worms when they were fed *S. maltophilia* (Fig. S3H). Our results suggest that the effect of *S. maltophilia* on *C. elegans* LD size was not dependent on food intake.

Next, we turned our attention to known lipid metabolic pathways in *C. elegans*. Wild type worms were fed OP50 or *S. maltophilia* from the L1 to the L4 stage and the relative expression level of selected metabolic genes was determined by real-time PCR. Our analysis clearly indicated that dietary *S. maltophilia* significantly increased the expression of *sbp-1, fat-5, fat-6* and *fat-7,* all of which are central of lipogenesis in *C. elegans* (Fig. 3A).

**Figure 3.**
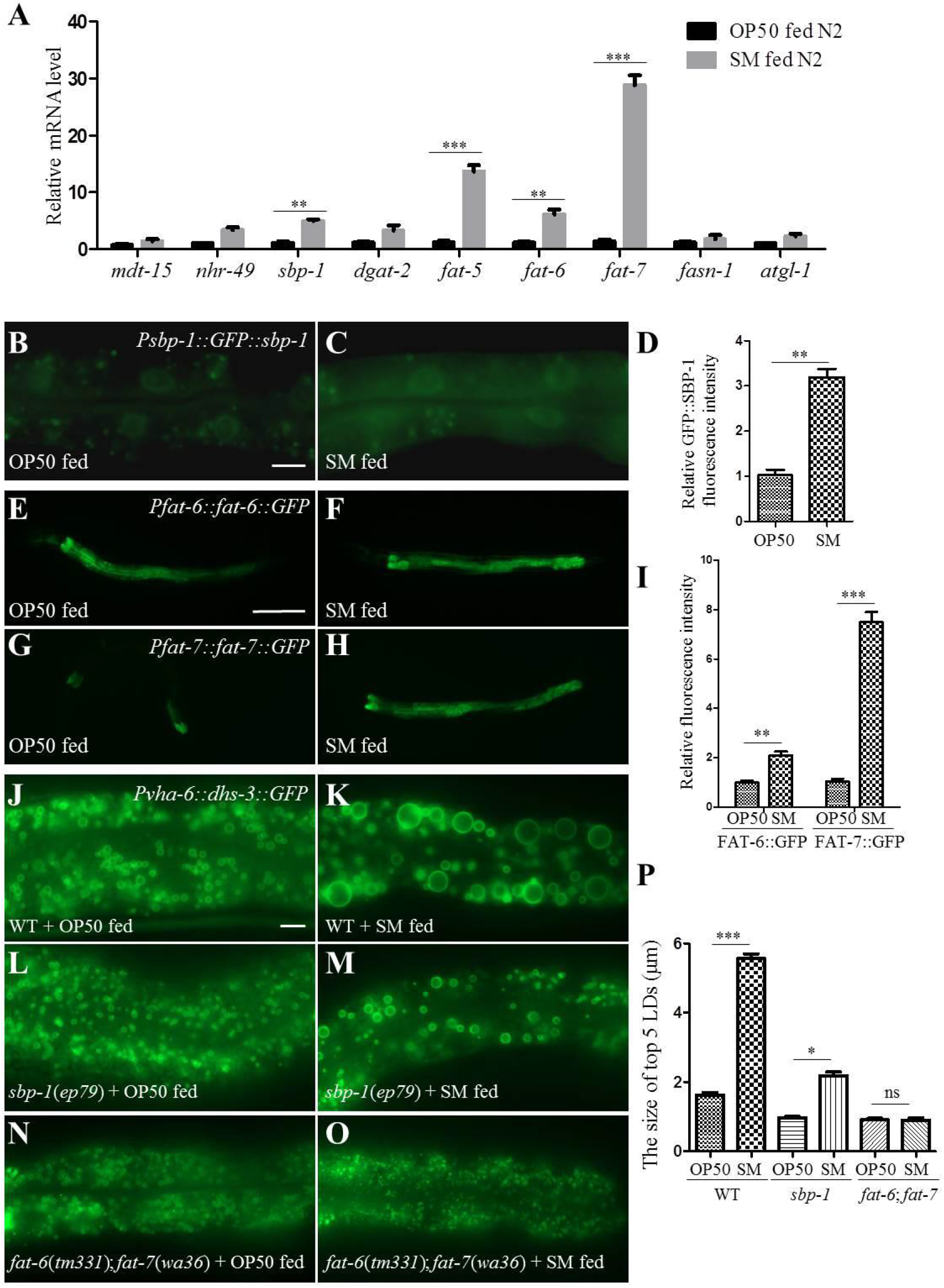
*Stenotrophomonas maltophilia* induces the LD enlargement in *C. elegans* through *sbp-1, fat-6; fat-7.* **(A)** RT-PCR result of the expression of lipid metabolism-related genes in OP50- and *S. maltophilia-.*fed N2 worms. Data represent mean ± SEM (n=3 for each independent experiment, \*\**P*<0.01, student *t*-test). **(B)** Fluorescence micrographs of GFP::SBP-1 in OP50-fed worms. Scale Bar, 5 μm. **(C)** Fluorescence micrographs of GFP::SBP-1 in *S. maltophilia-fed* worms. **(D)** Quantification of the GFP::SBP-1 fluorescence intensity in B and C. Data represent mean ± SEM (n=5 for each independent experiment, \*\**P*<0.01, student *t*-test). **(E)** Fluorescence micrographs of FAT-6::GFP in OP50-fed worms. Scale Bar, 100 μm. **(F)** Fluorescence micrographs of FAT-6::GFP in *S. maltophilia-fed* worms. **(G)** Fluorescence micrographs of FAT-7::GFP in OP50-fed worms. **(H)** Fluorescence micrographs of FAT-7::GFP in *S. maltophilia-fed* worms. **(I)** Quantification of the fluorescence intensity in E, F, G, and H. Data represent mean ± SEM (n=5 for each independent experiment, \*\**P*<0.01, student *t*-test). **(J)** Fluorescence micrographs of *Pvha-6::dhs-3::GFP* labeled LDs in the intestine of OP50-fed worms. Scale Bar, 5 μm. **(K)** Fluorescence micrographs of *Pvha-6::dhs-3::GFP* labeled LDs in *S. maltophilia-*fed worms. **(L)** Fluorescence micrographs of *Pvha-6::dhs-3::GFP* labeled LDs in *sbp-1(ep79)* mutant animals fed OP50. **(M)** Fluorescence micrographs of *Pvha-6::dhs-3::GFP* in *sbp-1(ep79)* mutant animals fed *S. maltophilia.* **(N)** Fluorescence micrographs of *Pvha-6::dhs-3::GFP* labeled LDs in *fat-6(tm331); fat-7(wa36)* double mutant animals fed OP50. **(O)** Fluorescence micrographs of *Pvha-6::dhs-3::GFP* labeled LDs in *fat-6(tm331); fat-7(wa36)* double mutant animals fed *S. maltophilia*. **(P)** Quantification of the LD diameter (J-O). Data represent mean ± SEM (n=5 for each independent experiment, \*\**P*<0.01, 0.01<\**P*<0.05, ns, no significance, student *t*-test).

SBP-1 is a basic helix-loop-helix (bHLH) transcription factor homologous to the mammalian sterol regulatory element binding proteins (SREBPs), which is required for lipid synthesis (Walker, Jacobs et al., 2011). FAT-6 and FAT-7 are acyl-CoA desaturases that function downstream of SBP-1 (Kim, Miyazaki et al., 2002, Miyazaki, Dobrzyn et al., 2004, Miyazaki, Flowers et al., 2007, Sampath, Miyazaki et al., 2007, Shi, Li et al., 2013, Yang, Vought et al., 2006). To verify the influence of *S. maltophilia* on these pathways, we fed *S. maltophilia* to *Psbp-1::GFP::sbp-1*, *Pfat-6::fat-6::GFP*, and *Pfat-7::fat-7::GFP* transgenic animals (Wu, Jiang et al., 2018). Feeding GFP::SBP-1 worms *S. maltophilia* resulted in robust induction of fluorescence in the whole worm (Fig. 3B, 3C and 3D). The expression of FAT-6::GFP and FAT-7::GFP was also increased with *S. maltophilia* feeding, compared to the OP50 control (Fig. 3E, 3F and 3I). We next determined the effect of *S. maltophilia* on *sbp-1(ep79)* mutant worms and *fat-6(tm331*); *fat-7(wa36)* double mutant worms. The ability of dietary *S. maltophilia* to increase LD size was blunted in *sbp-1*(*ep79*) animals (Fig. 3L, 3M, and 3P). Furthermore, dietary *S. maltophilia* had no effect on *fat-6(tm331); fat-7(wa36)* double mutant worms (Fig. 3N, 3O, and 3P). Taken together, we propose that the effect of *S. maltophilia* on host LDs was facilitated in part by the induction of the SBP-1/FAT-6/FAT-7 pathway.

### A Forward Genetic Screen for suppressors of *Stenotrophomonas maltophilia-induced* Lipid Droplet Expansion

We performed a forward genetic screen to identify *C. elegans* host factors that mediate the *S. maltophilia* effect on LDs. We used DHS-3::GFP or MDT-28::mCherry as LD markers. We screened 7,000 haploid genomes after chemical mutagenesis with ethyl-methane sulfonate (EMS), and isolated 7 mutant strains (Fig. 4A). Genetic mapping based on single nucleotide polymorphisms (SNPs) eventually led to the molecular cloning of three genes: *acs-13, dpy-9,* and *cyp-35B1* (Fig. 4D-J and S4). The *acs-13* gene encodes an ortholog of human ACSL1, 5, 6 (acyl-CoA synthetase long-chain family member 1, 5, 6). The *dpy-9* gene encodes a cuticular collagen family member with similarity to human collagen alpha 5, type IV. Finally, the *cyp-35B1* gene encodes a cytochrome P450 enzyme. Using complementation tests and RNAi, we confirmed that the loss of *acs-13*, *dpy-9*, or *cyp-35B1* function blocked the ability of *S. maltophilia* to induce LD expansion in *C. elegans* (Fig. 4D-4J).

**Figure 4.**
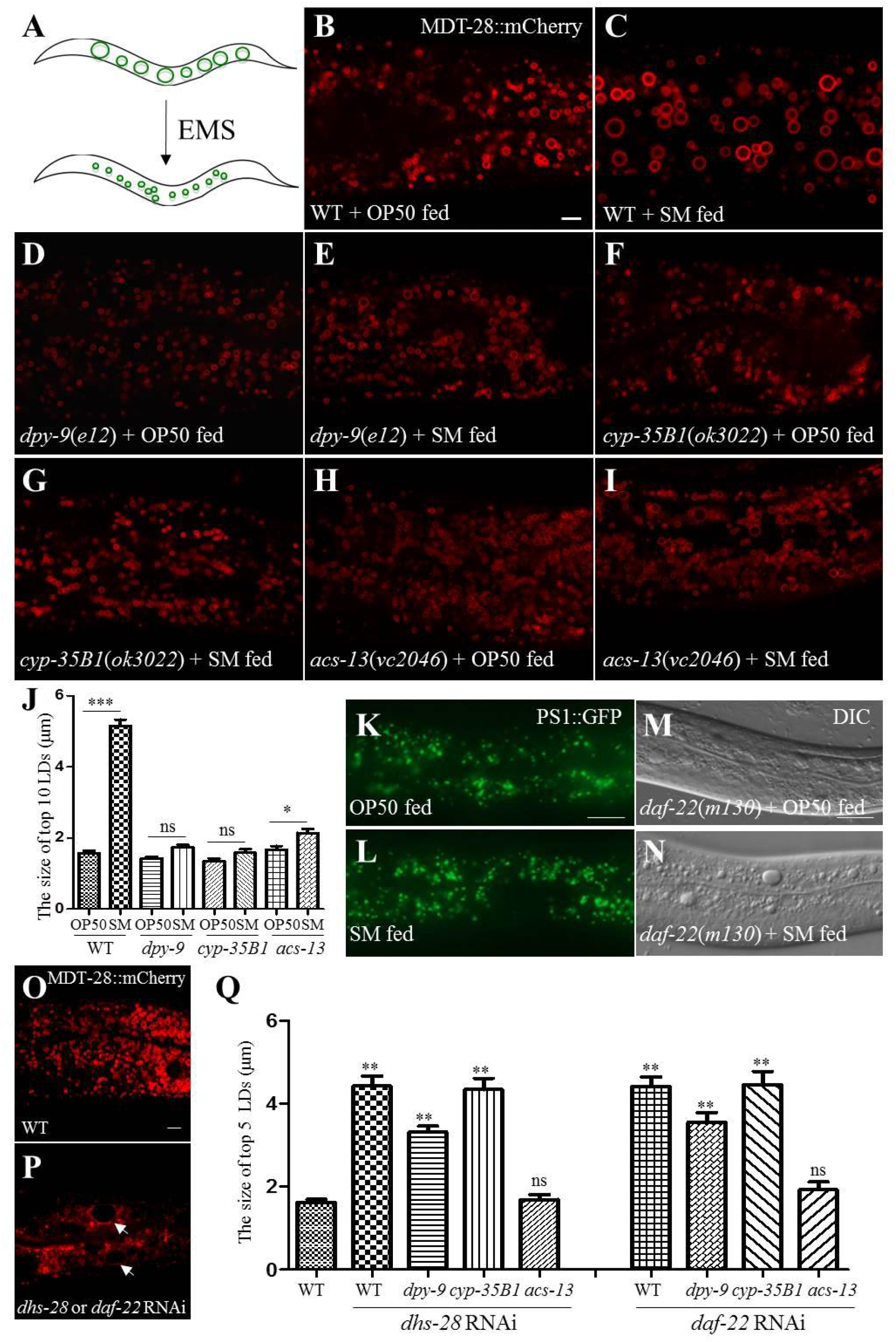
*dpy-9, cyp-35B1,* and *acs-13* suppress the *Stenotrophomonas maltophilia-induced* enlargement of LDs. **(A)** Schematic representation of the forward genetic screen. **(B)** Fluorescence micrographs of *Pmdt-28::mdt-28::mCherry* labeled LDs in OP50-fed worms. Scale Bar, 5 μm. **(C)** Fluorescence micrographs of *Pmdt-28::mdt-28::mCherry* labeled LDs in *S. maltophilia-fed* worms. **(D)** Fluorescence micrographs of *Pmdt-28::mdt-28::mCherry* labeled LDs in *dpy-9*(*e12*) mutant animals fed OP50. **(E)** Fluorescence micrographs of *Pmdt-28::mdt-28::mCherry* labeled LDs in *dpy-9*(*e12*) mutant animals fed *S. maltophilia*. **(F)** Fluorescence micrographs of *Pmdt-28::mdt-28::mCherry* labeled LDs in *cyp-35B1(ok3022)* mutant animals fed OP50. **(G)** Fluorescence micrographs of *Pmdt-28::mdt-28::mCherry* labeled LDs in *cyp-35B1(ok3022)* mutant animals fed *S. maltophilia*. **(H)** Fluorescence micrographs of *Pmdt-28::mdt-28::mCherry* labeled LDs in *acs-13(vc2046)* mutant animals fed OP50. **(I)** Fluorescence micrographs of *Pmdt-28::mdt-28::mCherry* labeled LDs in *acs-13(vc2046)* mutant animals fed *S. maltophilia.* **(J)** Quantification of LD diameter (B-I). Data represent mean ± SEM (n=5 for each independent experiment, \*\**P*<0.01, 0.01<\**P*<0.05, ns, no significance, student *t*-test). **(K)** Fluorescence micrographs of PS1::GFP labeled peroxisome in OP50-fed worms. Scale Bar, 5 μm. **(L)** Fluorescence micrographs of PS1::GFP labeled peroxisome in *S. maltophilia-fed* worms. **(M)** DIC images of LDs in *daf-22*(*m130*) mutant animals fed OP50. Scale Bar, 5 μm. **(N)** DIC images of LDs in *daf-22*(*m130*) mutant animals fed *S. maltophilia*. **(O)** Fluorescence micrographs of MDT-28::mCherry labeled LDs in OP50-fed worms. Scale Bar, 5 μm. **(P)** Fluorescence micrographs of MDT-28::mCherry labeled LDs in *dhs-28* or *daf-22* RNAi animals. The white arrows point to the enlarged LDs **(Q)** Quantification of LD diameter (O and P). Data represent mean ± SEM (n=5 for each independent experiment, \*\**P*<0.01, ns, no significance, two-way ANOVA).

We next investigated if loss of *acs-13*, *dpy-9*, or *cyp-35B1* function could normalize fat storage in *C. elegans* mutants that are known to accumulate excess fat. Notably, mutations in *dhs-28* and *daf-22* are known to impair peroxisomal β-oxidation and induce LD expansion in *C. elegans* (Butcher, Ragains et al., 2009, Zhang, Box et al., 2010). We analyzed OP50- and *S. maltophilia*- fed worms and found no significant difference in peroxisome morphology (Fig. 4K and 4L). Feeding *daf-22(m130)* mutants *S. maltophilia* resulted in a significant increase in the number and size of LDs (Fig. 4M and 4N), suggesting that dietary *S. maltophilia* acted in parallel of the peroxisomal β-oxidation pathway, which is responsible for fat catabolism. Finally, we recapitulated the LD expansion phenotype by knocking down *dhs-28* and *daf-22* by RNAi in wild type worms (Fig. 4O and 4P). Next, we knocked down *dhs-28* and *daf-22* in *acs-13*, *dpy-9*, and *cyp-35B1* mutant worms. The *acs-13* mutation suppressed the formation of enlarged LDs in *dhs-28* or *daf-22* deficient worms. In contrast, mutations in *dpy-9* or *cyp-35B1* had no effect (Fig. 4Q). Therefore, we concluded that CYP-35B1 and DPY-9 are specific host factors that are required for dietary *S. maltophilia* to promote LD expansion.

### *Stenotrophomonas maltophilia* Induced Lipid Droplet expansion by enhancing ER-LD interaction

The ER is the primary site where TAG is synthesized (Fagone & Jackowski, 2009, Sorger & Daum, 2003). Therefore, we investigated the effects of *S. maltophilia* feeding on the ER. We used two reporters, *Phyp-7::TRAM-1::GFP* and *Pvha-6::SEL-1(1-79)::mCherry::HDEL,* to mark the ER membrane and lumen, respectively. After 2 days of *S. maltophilia* feeding, ring-shaped structures were observed in the hypodermis of the TRAM-1::GFP worms (Fig. S5C-5F). We also observed ER-wrapped LDs in the intestine of the mCherry::HDEL worms (Fig. S5G-5L). It is plausible that the remodeling of the ER morphology upon *S. maltophilia* feeding supports LD expansion.

We further investigated the ER-LD interaction using electron microscopy. In worms fed the *E. coli* OP50 diet, few connections between the ER and LDs were observed (Fig. 5A and 5D). In contrast, in worms fed the *S. maltophilia* diet, we readily noted LDs that were connected to the ER through small tubular structures (Fig. 5B, 5C, 5E, and 5F). To observe the contact sites in detail, electron tomography was performed. The images demonstrate that the connection between LDs and the ER was hollow (Fig. 5G–5J). Similar structures have previously been reported (Prinz, 2013). We name this structure ER-LD bridge.

**Figure 5.**
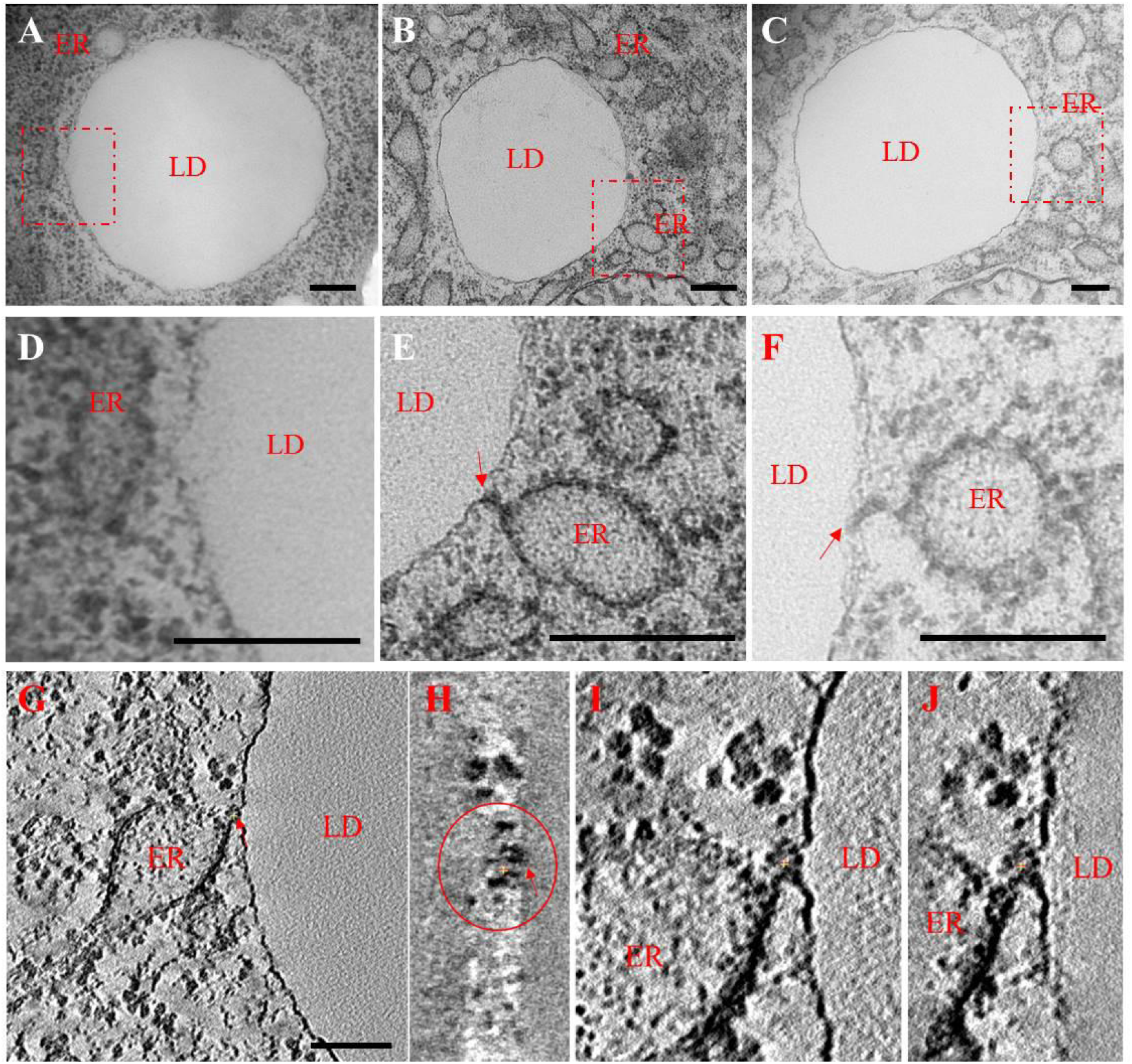
*Stenotrophomonas maltophilia* induces the formation of ER-LD bridge. **(A)** Morphology of ER and LD in N2 animals fed OP50. Scale Bar, 200 nm. **(B)** Morphology of ER and LD in N2 fed *S. maltophilia*. Scale Bar, 200 nm. **(C)** As in (B), the other results for the morphology of ER and LD in N2 fed *S. maltophilia.* Scale Bar, 200 nm. **(D, E, and F)** Insets enlarged for A, B, and C. Scale Bar, 200 nm **(G)** The ER-LD bridge in N2 fed *S. maltophilia* displayed using the tomography method. Scale Bar, 100 nm. **(H)** Inset enlarged for G. **(I and J)** A different consecutive section and its enlarged inset.

### *Stenotrophomonas maltophilia* Induced ER-LD Contacts

To further understand how *S. maltophilia*-induced LD expansion was instigated by ER remodeling, we examined the localization of SEIP-1, which is highly enriched in an ER subdomain that associates tightly with LDs (Cao, Hao et al., 2019). SEIP-1 is the *C. elegans* ortholog of seipin, which functions at ER-LD contact sites (Grippa, Buxo et al., 2015, Pagac, Cooper et al., 2016, Salo, Belevich et al., 2016, Wang, Becuwe et al., 2016). To visualize SEIP-1 positive ER subdomain (peri-LD cages), we used two reporter strains that expressed SEIP-1::GFP fusion proteins either ubiquitously *(hjSi189)* or specifically in the intestine *(hjSi3)* (Cao et al., 2019).

When *hjSi189* worms were fed *E. coli* OP50, few intestinal LDs were associated with SEIP-1::GFP cages (Fig. 6B and 6C). In contrast, almost all intestinal LDs were enwrapped by SEIP-1::GFP cages when *hjSi189* transgenic worms were fed *S. maltophilia*, coincident of LD expansion (Fig. 6E and 6F). To investigate further, we conducted 3D structured illumination microscopy (3D-SIM) on *S. maltophilia*- and OP50-fed *hjSi189* (Fig. 7A-C, S6). We found that more than 95% of LDs were associated with SEIP-1::GFP positive structures in worms on the *S. maltophilia* diet. We also quantified the fraction of biochemically purified LDs that were decorated with SEIP-1::GFP. LDs were isolated from *hjSi189* worms fed OP50 or *S. maltophilia* for 2.5 days. Again, few LDs were found to retain SEIP-1::GFP in the OP50-fed group while 100% of LDs from the *S. maltophilia-fed* group showed SEIP-1::GFP signals (Fig. 6G–6M). These results are consistent with the model that the SEIP-1 positive ER subdomain promotes LD expansion, and that *S. maltophilia* feeding significantly increases the coverage of such subdomain, thereby supporting widespread LD expansion.

**Figure 6.**
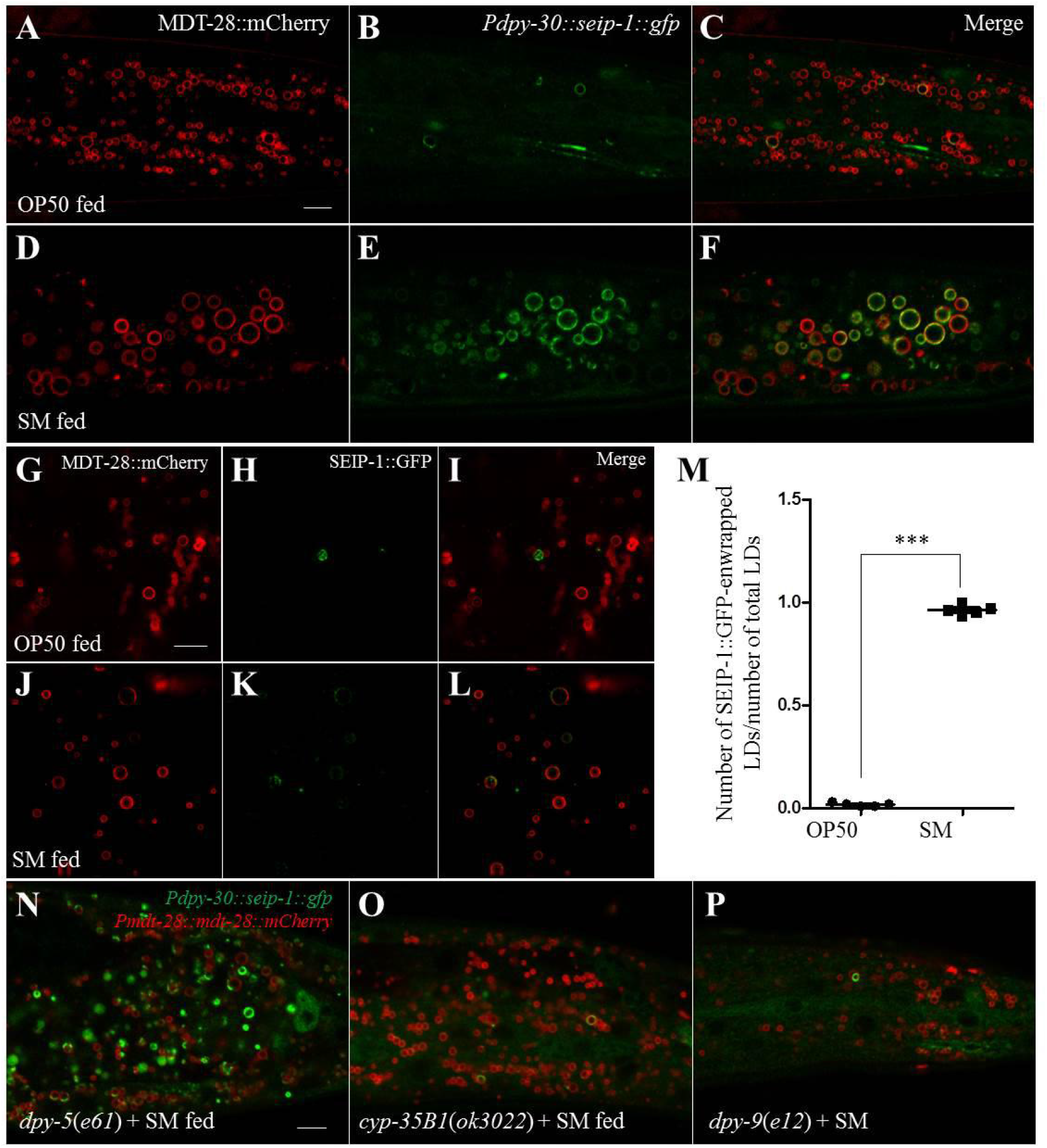
*Stenotrophomonas maltophilia* induces SEIP-1::GFP-labeled ER enwrapping LDs. **(A)** Fluorescence micrographs of *Pmdt-28::mdt-28::mCherry* labeled LDs in OP50-fed worms. Scale Bar, 5 μm. **(B)** Fluorescence micrographs of *Pdpy-30::seip-1::gfp* labeled ER in OP50-fed worms. **(C)** As in (A), but with (B) merged. **(D)** Fluorescence micrographs of *Pmdt-28::mdt-28::mCherry* labeled LDs in *S. maltophilia-fed* worms. **(E)** Fluorescence micrographs of *Pdpy-30::seip-1::gfp* labeled ER in *S. maltophilia-fed* worms. **(F)** As in (D), but with (E) merged. **(G)** Fluorescence micrographs of LDs purified from transgenic animal expressing MDT-28::mCherry. Scale Bar, 5 μm. **(H)** Fluorescence micrographs of LDs purified from OP50-fed transgenic animal expressing *Pdpy-30::seip-1::gfp*. **(I)** as in (G), but with gfp and mCherry signals merged. **(J)** Fluorescence micrographs of LDs purified from *S. maltophilia*-fed transgenic animal expressing MDT-28::mCherry. **(K)** Fluorescence micrographs of LDs purified from *S. maltophilia*-fed transgenic animal expressing *Pdpy-30::seip-1::gfp*. **(L)** as in (J), but with gfp and mCherry signals merged. **(M)** Quantification of ratio of the number of seip-1::GFP positive LDs to number of total LDs. Data represent mean ± SEM (n=5 for each independent experiment, \*\**P*<0.01, student *t*-test). **(N)** Fluorescence micrographs of *Pmdt-28::mdt-28::mCherry* and *Pdpy-30::seip-1::gfp* in *dpy-5(e61)* mutant animals. Scale Bar, 5 μm. **(O)** Fluorescence micrographs of *Pmdt-28::mdt-28::mCherry* and *Pdpy-30::seip-1::gfp* in *dpy-9(e12)* mutant animals. **(P)** Fluorescence micrographs of *Pmdt-28::mdt-28::mCherry* and *Pdpy-30::seip-1::gfp* in *cyp-35B1(ok3022)* mutant animals.

Next, we investigated if loss of CYP-35B1 and DPY-9 function could block *S. maltophilia*-induced LD expansion by interfering with SEIP-1 enrichment in peri-LD cages. Accordingly, we introduced *cyp-35B1*, *dpy-9*, and *dyp-5* (negative control) mutations into the SEIP-1::GFP reporter strain *hjSi189.* After feeding *S. maltophilia* for 2 days, no SEIP-1::GFP positive, peri-LD cages were observed in *cyp-35B1* and *dpy-9* mutant animals (Fig. 6N, 6O and 6P). It is conceivable that CYP-35B1 and DPY-9 directly or indirectly act in pathways that remodel the membrane environment of peri-LD cages. Such remodeling is crucial for the recruitment of SEIP-1 and additional proteins that are critical for LD expansion.

### Requirement of Proteins Involved in ER-LD Interactions for *Stenotrophomonas maltophilia*-induced LD expansion

To determine if SEIP-1 is important for LD expansion induced by dietary *S. maltophilia,* we fed *S. maltophilia* to *seip-1(tm4221)* mutant animals (Fig. 7G). Quantification revealed that loss of SEIP-1 function only partially suppressed the effect of *S. maltophilia* feeding on LDs (Fig. 7H). Our results hinted at additional proteins that may reside in ER-LD contacts that mediate the effect of dietary *S. maltophilia*. To this end, we investigated *rab-18* and *dgat-2* since they have been reported to be involved in ER-LD interaction (Jayson, Arlt et al., 2018, Jin, McFie et al., 2014, Li, Luo et al., 2017, Xu, Li et al., 2018, Xu, Zhang et al., 2012). We examined the effect of *rab-18*, and *dgat-2* mutations on the size of LDs in *C. elegans* after *S. maltophilia* feeding. As shown in Figure 8I–8M, loss of DGAT-2 function partially suppressed LD expansion. However, loss of RAB-18 function had no effect in this experimental context.

**Figure 7.**
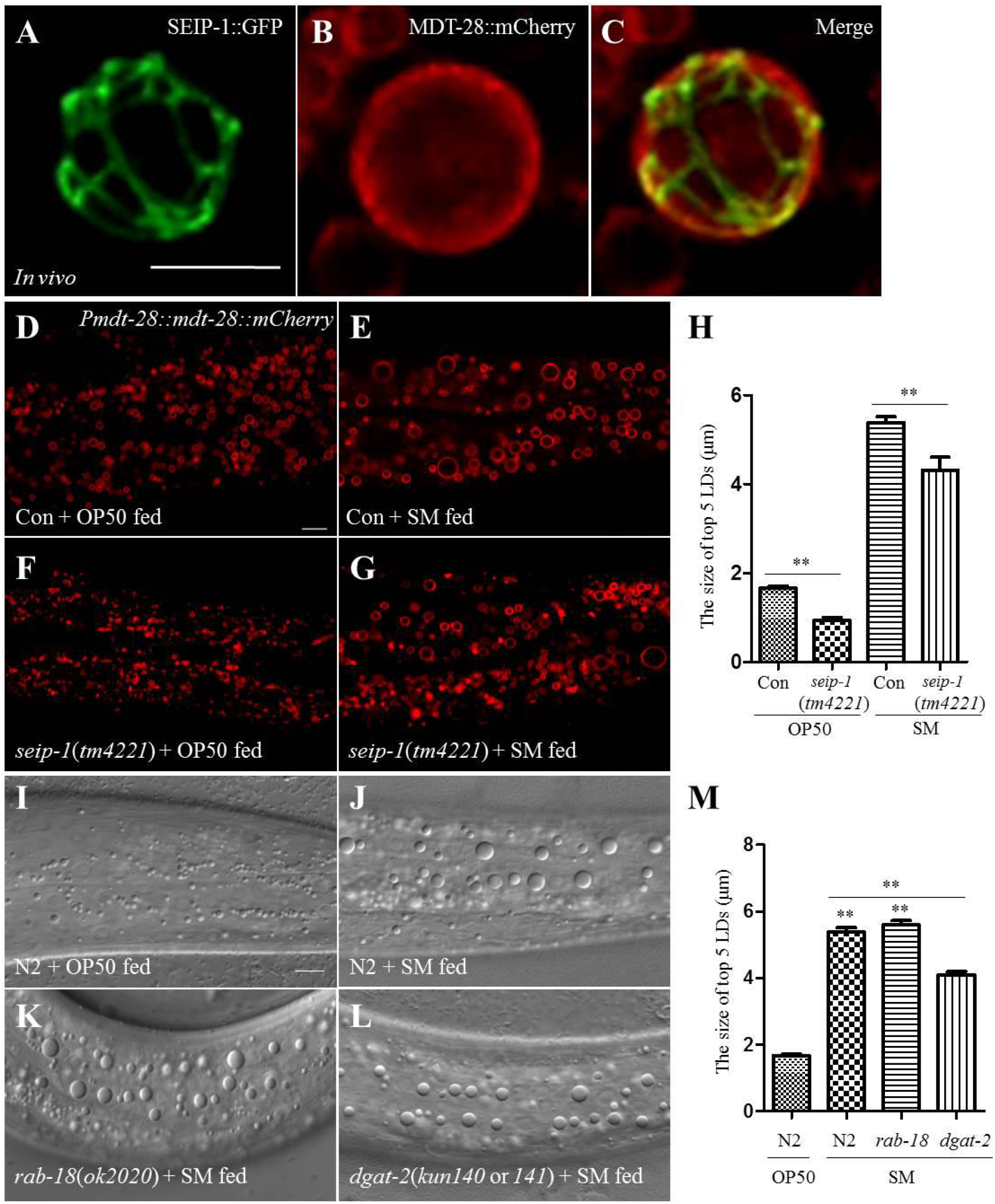
Proteins involved in ER-LD interaction partially suppress the *Stenotrophomonas maltophilia-*enlarged LDs. **(A) 3D-**SIM images of SEIP-1::GFP labeled ER in OP50-fed worms. Scale Bar, 1 μm. **(B) 3D-**SIM images of MDT-28::mCherry labeled LDs in OP50-fed worms. **(C)** As in (A), but with (B) merged. **(D)** Fluorescence micrographs of *Pmdt-28::mdt-28::mCherry* labeled LDs in OP50-fed worms. Scale Bar, 5 μm. **(E)** Fluorescence micrographs of *Pmdt-28::mdt-28::mCherry* labeled LDs in *S. maltophilia-fed* worms. **(F)** Fluorescence micrographs of *Pmdt-28::mdt-28::mCherry* labeled LDs in *seip-1(tm4221)* mutant animals fed OP50. **(G)** Fluorescence micrographs of *Pmdt-28::mdt-28::mCherry* labeled LDs in *seip-1(tm4221)* mutant animals fed *S. maltophilia.* **(H)** Quantification of the LD diameter for (D-G). Data represent mean ± SEM (n=5 for each independent experiment, \*\**P*<0.01, student *t*-test). **(I)** DIC images of LDs in OP50-fed worms. Scale Bar, 5 μm. **(J)** DIC images of LDs in worms fed *S. maltophilia*. **(K)** DIC images of LDs in *rab-18(ok2020)* mutant animals fed *S. maltophilia.* **(L)** DIC images of LDs in *dgat-2(kun140* or *141*) mutant animals fed *S. maltophilia*. **(M)** Quantification of LD diameter for (J-M). Data represent mean ± SEM (n=5 for each independent experiment, \*\**P*<0.01, two-way ANOVA).

## Discussion

In this paper, we show that dietary *S. maltophilia* promoted organismal fat storage in *C. elegans* through a lipogenic transcriptional program that consists of SBP-1, FAT-6 and FAT-7. In addition, *S. maltophilia* feeding enhanced CYP-35B1-, DPY-9-dependent ER-LD interaction in the intestine and hypodermis, which was prerequisite for LD expansion. Collectively, we hypothesize that the increased contact, via ER-LD bridges, facilitates the transfer of lipid from the ER to LDs (Fig. 8).

**Figure 8.**
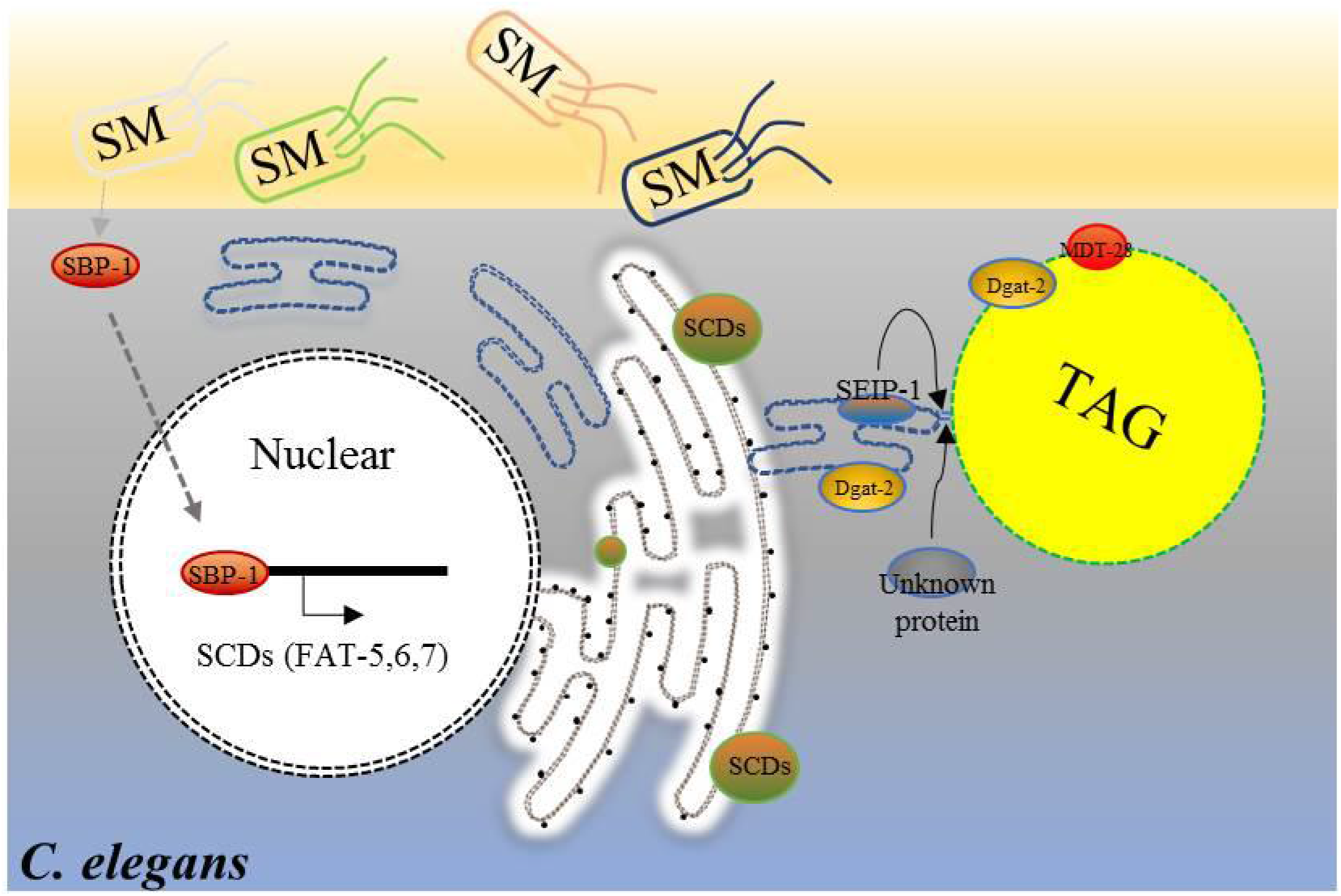
Model for the enlargement of LDs in *C. elegans* fed on *S. maltophilia.* Different bacteria have different effects on the obesity of nematodes. Compared with the standard *E. coli* OP50, *S. maltophilia* induces the accumulation of enlarged LDs through an increase in lipogenesis by up-regulating genes *(sbp-1/fat-6; fat-7),* and increasing the interaction between ER and LDs in nematodes. The ER-LD bridge likely facilitates the transfer of lipid from ER to LDs in *C. elegans*.

The use of *C. elegans* to study host-pathogen interaction was pioneered by the Ausubel lab, which stemmed from the observations that *Pseudomonas Aeruginosa* PA14 could effectively kill worms (Tan, Mahajan-Miklos et al., 1999). *C. elegans* genetic and molecular analyses led to the discovery of conserved innate immune response pathways that counter *P. Aeruginosa* infection (Irazoqui & Ausubel, 2010, Kim, Feinbaum et al., 2002). The systematic response appears to require environmental sensing and inter-tissue communication between the nervous system, hypodermis and intestine (Meisel & Kim, 2014, Singh & Aballay, 2019, Wani, Goswamy et al., 2019). Ultimately, multiple transcriptional networks coordinately mount gene expression programs that support the defense against the pathogen (Fletcher, Tillman et al., 2019, Troemel, Chu et al., 2006). Besides *P. Aeruginosa,* many other pathogenic bacteria, such as *E. faecalis, S. aureus, S. typhimurium* and *C. neoformans*, have subsequently been shown to effectively kill worms (Kim & Ewbank, 2018). However, relatively less is known about the impact of pathogenic bacteria on *C. elegans* physiology during chronic infection. Here, we discovered that environmental isolates of the pathogenic bacterium *S. maltophilia* did not kill worms. Instead, worms maintained on a *S. maltophilia* diet grew slower and accumulated excess neutral fat. Accordingly, the p38 MAP kinase pathway, which is central to innate immunity against lethal pathogens in *C. elegans*, did not play a role in the induction of fat storage in response to dietary *S. maltophilia*. Our results indicate that *S. maltophilia* may exert metabolic burden to its host. Further investigation is needed to elucidate the casual factors from *S. maltophilia* that modulate host metabolism.

Since *S. maltophilia* is commonly found in immunocompromised, hospitalized patients (Crossman, Gould et al., 2008), there is an urgent need to understand the host response to this opportunistic pathogen, in order to identify cellular proteins that may serve as therapeutic targets. From a forward genetic screen, we found that loss of function mutations in *cyp-35B1* and *dpy-9* rendered the worms resistant to *S. maltophilia* induced fat accumulation. The *cyp-35B1* gene encodes a cytochrome P450 enzyme, and was originally identified as a transcriptional target of the DAF-2 insulin/IGF-I-like signaling pathway (Iser, Wilson et al., 2011, Murphy, McCarroll et al., 2003). Reporter assays suggested that it is expressed in the intestine (Iser et al., 2011). In contrast, the *dpy-9* gene encodes an extracellular matrix collagen that is presumably expressed in the hypodermis. The lack of *dpy-9* compromises cuticle integrity that may give rise to osmotic stress (Rohlfing, Miteva et al., 2010). Based on the functional annotation of these two proteins, it is currently unclear how the loss of CYP-35B1 or DPY-9 function in distinct tissues confers resistance to *S. maltophilia*. It is plausible that CYP-35B1 and DPY-9 are part of a wider genetic network that facilitates inter-tissue communication in response to *S. maltophilia*. Although further analysis is needed to identify other components of this network, our results clearly suggest that one of its outputs is required to sustain ER-LD contacts, which in turn support fat accumulation in LDs.

In summary, our work has established a new host-microbe experimental paradigm. Chronic exposure of *C. elegans* to *S. maltophlia* reproducibly caused metabolic remodeling in multiple tissues. Future experiments will be directed to studying microbial factors that are responsible for such remodeling via CYP-35B1, DPY-9 and additional yet-to-be identified host factors.

## Materials and Methods

### Nematode strains and growth conditions

All *C. elegans* strains were handled and maintained following standard procedures (Brenner, 1974). The N2 Bristol strain, *dpy-9(e12)* and *dpy-5(e61)* were obtained from the *Caenorhabditis* Genetics Center. *sbp-1*(*ep79*), *fat-6*(*tm331*), *fat-7(wa36), epEx307[Psbp-1::GFP::sbp-1], [Pfat-7::fat-7::GFP]*, and *[Pfat-6::fat-6::GFP]* were gifts from Bin Liang’s laboratory. *seip-1(tm4211)* was obtained from Xun Huang’s laboratory. *hjSi158[vha-6p::SEL-1(1-79)::mCherry::HDEL::let-858 3’ UTR], hjSi3[vha-6p::seip-1 cDNA::GFP_TEV_3xFLAG::let-858 3’ UTR], ldrIs3[vha-6p::dhs-3::gfp]*, and *hjSi189[dpy-30p::seip-1 cDNA::GFP_TEV_3xFLAG::tbb-2 3’ UTR]* were from Ho Yi Mak’s laboratory. *sek-1*(*km4*), *pmk-1*(*ku54*), *Phyp-7::tram-1::gfp,* and *Phyp-7::PS1::gfp* were from Hong Zhang’s laboratory. *ldrIs2[mdt-28p::mdt-28::mCherry, unc-76(+)]* strains were constructed in our laboratory. All experimental animals were maintained at 20°C.

*S. maltophilia* and OP50 were cultured on LB and NGM plates.

### Forward genetic screen and mutant mapping

To screen the mutant animals with a suppressed phenotype of LD induced by *S. maltophilia*, *ldrIs3* was mutagenized with ethyl methane sulfonate (EMS) as previously described (Encalada, Martin et al., 2000, Kemphues, Kusch et al., 1988). 7,000 haploid genomes were screened and mutant phenotype in the F2 worms were selected to observe the LD phenotype. Mutant worms were backcrossed with *ldrIs3* at least four times and their LD phenotypes were studied using a ZEISS Imager M2. The stable mutant animals were mapped using single-nucleotide polymorphism (SNP) mapping (Davis, Hammarlund et al., 2005).

### Isolation of lipid droplets

LDs were isolated using a method described previously (Ding, Zhang et al., 2013, Zhang et al., 2012). Briefly, synchronized L1 worms were cultured on NGM plates and the L4 larval stage worms were collected in M9 buffer (22 mM KH_2_PO_4_, 42 mM Na_2_HPO_4_, 86 mM NaCl, 1 M MgSO_4_). The worms were washed three times in Buffer A (25 mM Tricine, pH 7.6, 250 mM Sucrose, and 0.2 mM phenylmethylsulfonyl fluoride) and were then homogenized to obtain whole body lysates. The lysates were centrifuged at different speeds (4,000 rpm, 6,000 rpm and 10,000 rpm) in SW40 tubes to obtain LDs of different diameters. The LD fraction was collected from the top layer and was washed three times with Buffer B (20 mM HEPES, 100 mM KCl, 2 mM MgCl2, pH 7.4). Images of the isolated LDs were obtained using a ZEISS Imager M2 or LSM880.

### Measurement of triacylglycerol

Synchronized L4 worms were washed three times with M9, and were dissolved in 200 μl 1% Triton X-100 by sonication. Then, the worm lysates were centrifuged at 12,000 rpm for 3 min. The triacylglycerol (TAG) content of the supernatants was measured using the Triacylglycerol Assay Kit (Liu et al., 2018). The Pierce BCA Protein Assay Kit (Thermo, USA) was used to quantify the proteins (Liu et al., 2018).

### Lipid extraction and analysis

Bacterial lipid extraction, separation, and analysis were conducted as described previously (Shi et al., 2013). Bacteria cultured under standard culture conditions were collected into glass tubes and water was removed with a Pasteur pipet. To each bacterial pellet 1 ml of MeOH and 2.5% H_2_SO_4_ was added. Fatty acids were extracted and converted into methyl esters by heating at 70°C for 60 minutes. Then the extractions were incubated at 25°C for 5 min followed by addition of 0.2 ml of hexane and 1.5 ml of water. The mixtures were shaken vigorously and then centrifuged 10,000 rpm for 1 minute in a clinical centrifuge. The fatty acid methyl esters contained in the top hexane-rich fraction were collected and 2 μl were analyzed using an Agilent 6890 series gas chromatograph equipped with a 20 m x 0.25 mm SP-2380 column (Supelco, Bellefonte, PA) and a flame ionization detector. The Gas Chromatography (GC) was programmed for an initial temperature of 120°C for 1 min followed by an increase of 10°C per minute up to 190°C followed by an increase of 2°C per minute to 200°C. Peak identity was determined by mass spectroscopy.

### Transposon-based Forward Genetic Screen

The *S. maltophilia* transposon insertion mutant library was constructed using an EZ::TN <KAN-2> Tnp transposome kit (Epicentre) as described previously and in accordance with the manufacturer’s protocol (Weisenfeld, Kumar et al., 2017). The kanamycin-resistant transformants were selected and fed the worms. To identify mutants which did not result in enlarged LDs when fed to *C. elegans*.

### Fluorescence imaging of *C. elegans*

Worms were immobilized with 0.5 mg/ml levamisole in M9 buffer and then transferred to a glass slide. Fluorescence images of L4 larval animals were obtained using a laser scanning confocal microscope (LSM 710 Meta, LSM880 Meta, ZEISS). The imaging position is the same as the nematode’s tail. Images were processed and viewed using ZEN 2011 software (ZEISS).

### Quantitative RT-PCR analysis

Total RNA was isolated using Trizol reagent (Tawe, Eschbach et al., 1998). Moloney murine leukemia virus (M-MuLV) reverse transcriptase with random hexamer primers was used to synthesize the cDNA. RT-PCR was performed on a CFX96 real-time system with SYBR green. Relative expression levels of all mRNAs were normalized to *ama-1* mRNA.

### Behavioral experiment

OP50 and *S. maltophilia* were cultured and dropped on NGM plates. The bacteria were allowed to grow for 24 hours. The synchronized L4 larval nematodes were placed equidistant between the OP50 and *S. maltophilia* cultures. After 12 h, the number of nematodes on the bacterial colonies was determined.

### RNAi assay of *C. elegans*

L4440 was used as the control for the RNAi assay. The RNAi for *dhs-28* and *daf-22* were from the Ahringer RNAi library. The synchronized L1 worms were cultured on the RNAi NGM plates at 20°C to generate F4 worms for phenotypic analysis.

### Brood size analysis

Approximately 20 L4 worms were removed from synchronized mothers, fed with OP50, and transferred to NGM plates seeded with the appropriate bacteria, in triplicate. The worms were transferred to new plates every 24 hours three times until no additional embryos were produced. The number of embryos was counted every 24 hours (Brock, Browse et al., 2007, Brooks et al., 2009).

### Growth rate

Two-day old adults fed with the relevant bacteria were treated with hypochlorite to obtain embryos. The embryos were plated on NGM plates (~100/plate) seeded with different bacteria. The percentage of L4 worms out of the total worms was determined at various time points (Liu et al., 2018).

### Pharyngeal pumping measurements

L4 stage worms were picked from NGM plates. After 24 hours, the pumping rate on different food was recorded. Worms were viewed under an optical microscope and pumping rates were measured by visual observation. One pharyngeal pump was defined as a complete forward and backward movement of the grinder in the pharynx.

The rate was counted for a period of 30 sec (Raizen, Lee et al., 1995).

### Electron microscopy assay

Samples were prepared using previously described methods (Li, Ji et al., 2017). 2D images were acquired using transmission electron microscope (FEI Tecnai Sprit 120 kV). 3D images were acquired using transmission electron microscope (FEI Tecnai Sprit 120 kV and Serial EM software).

### Data analysis

All numerical data were plotted as mean ± SEM unless otherwise indicated. The statistical analysis was performed using GraphPad Prism 5 and Image J (NIH, USA). For the number of samples, please refer to the figure legend of each experiment. Determination of significance between groups was performed using Student *t*-tests, one-way ANOVA or two-way ANOVA as indicated.

## Acknowledgements

The authors thank Dr. John Zehmer for his critical reading and useful suggestions and Ms. Shuoguo Li for her help in taking and analyzing the images. The authors also thank the *Caenorhabditis* Genome Center (CGC) and National BioResource Project (NBRP) for providing strains. This work was supported by the National Key R&D Program of China (Grant No. 2016YFA0500100, 2018YFA0800700 and 2018YFA0800900), National Natural Science Foundation of China (Grant No. 91857201, 91954108, 31571388, 31671402, 31671233, 31701018 and U1702288). This work was also supported by the “Personalized Medicines Molecular Signature-based Drug Discovery and Development”, Strategic Priority Research Program of the Chinese Academy of Sciences, Grant No. XDA12040218. This work was also supported by the CAS-Croucher Joint Laboratory Project, Project No. CAS16SC01.

## Author Contributions

P.L., H.Y.M., and H.Z. conceived the project. K.X., Y.L., and X.L. carried out experiments and data analysis. Manuscript was written by K.X., S.Z., H.Y.M. and P.L.

