## Supplemental files for "Dietary *S. maltophilia* promotes fat storage by enhancing lipogenesis and ER-LD contacts in *C. elegans*"

**Supporting Information**
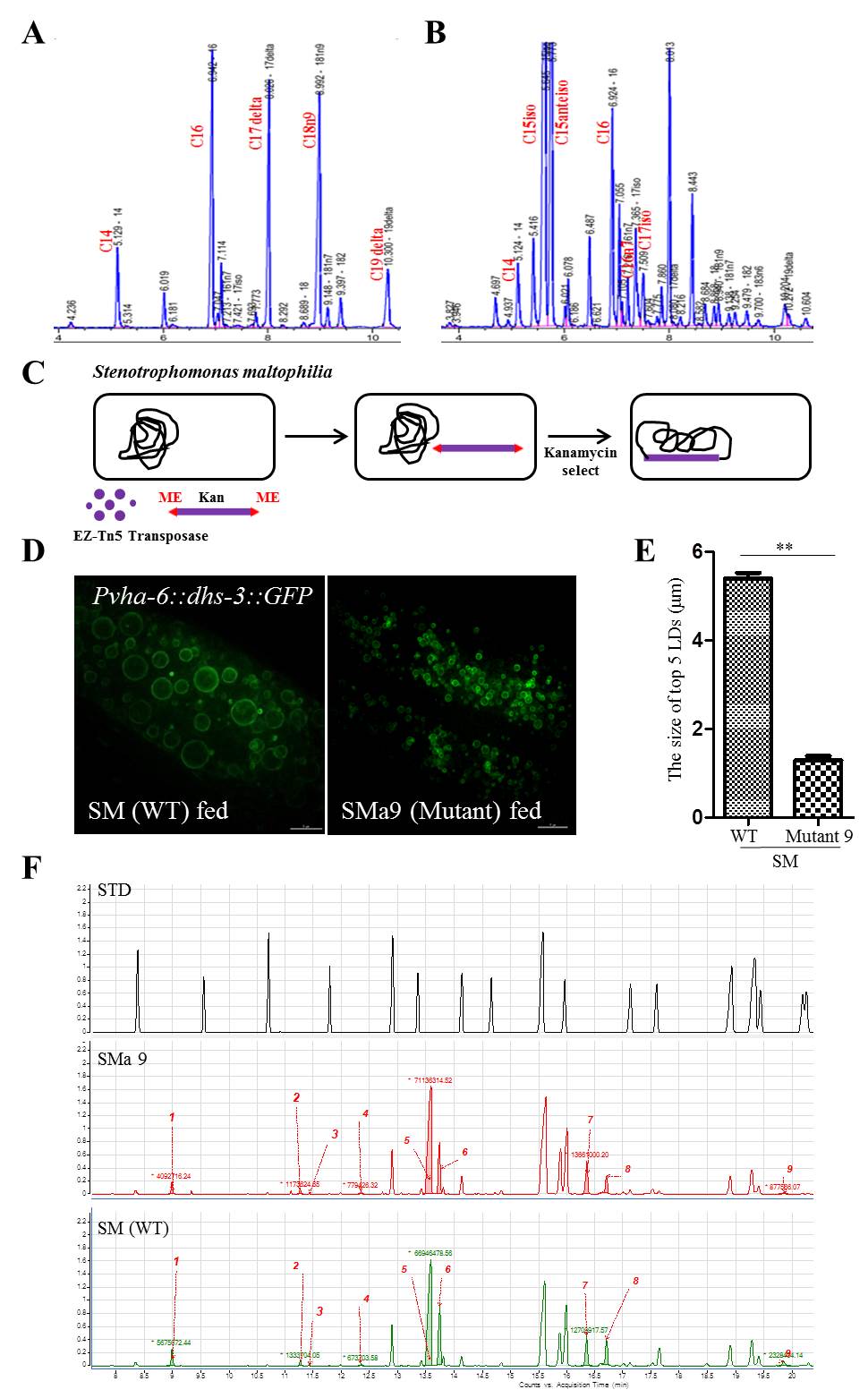

**Figure S1 Differential fatty acids were not the cause for the enlargement of LDs in *S. maltophilia*-fed nematode.**

**(A)** Fatty acid composition of *E. coli* strain. **(B)** Fatty acid composition of *S. maltophilia* strain. **(C)** The cartoon to show the method to obtain the bacterial mutants. **(D)** Fluorescence micrographs of *Pvha-6::dhs-3::GFP* in *S. maltophilia* (WT) and *S. maltophilia* (SMa9)*-*fed worms. **(E)** Quantification of LD diameter (D). Data represent mean ± SEM (n=5 for each independent experiment, ***P*<0.01, student *t*-test). **(F)** Fatty acid composition of *S. maltophilia* (WT) and *S. maltophilia* (SMa9) strains. Individual peaks are as follows: 1. Methyl 8-methyl-decanoate, 2. Methyl 11-methyl-dodecanoate, 3. Dodecanoic acid, 4. Tridecanoic acid, 5,6 Methyl 13-methyltetradecanoate, 7. Hexadecanoic acid, 8. 11-Hexadecenoic acid, 9. Octadecanoic acid.

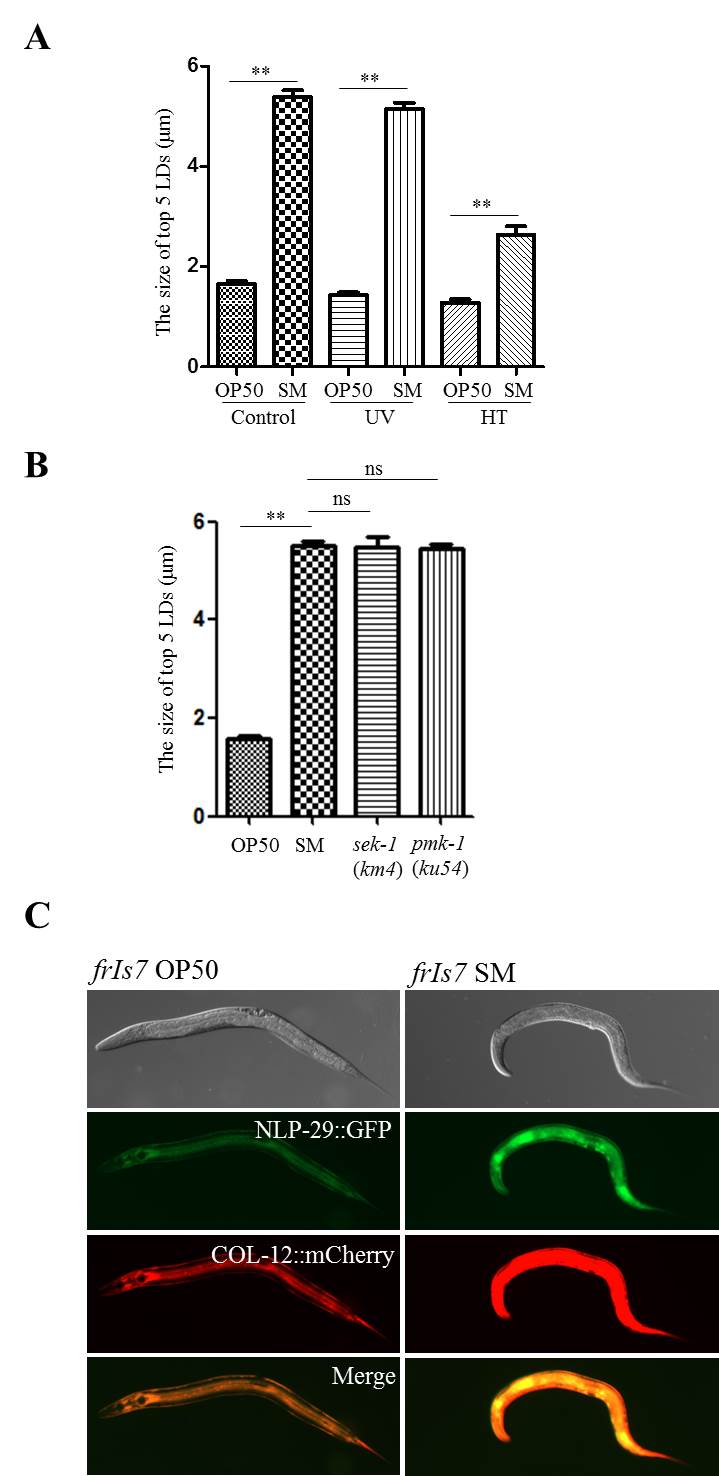

**Figure S2 *Stenotrophomonas maltophilia* increases NLP-29 protein level in *C. elegans.***

**(A**) Quantification of the LD diameter in N2 animals after feeding with untreated*,* UV-killed*,* high temperature-killed OP50 and *S. maltophilia*. (n=5 for each independent experiment, student *t*-test). **(B)** Quantification of the LD diameter in *sek-1*(*km4*) and *pmk-1*(*ku54*) animals after feeding with *S. maltophilia*. (n=5 for each independent experiment, student *t*-test). **(C)** *frIs7* strain worms were fed with OP50 or *S. maltophilia* from L1 to L4 stage. The protein expression level is shown by fluorescence images. Scale Bar, 100 μm.

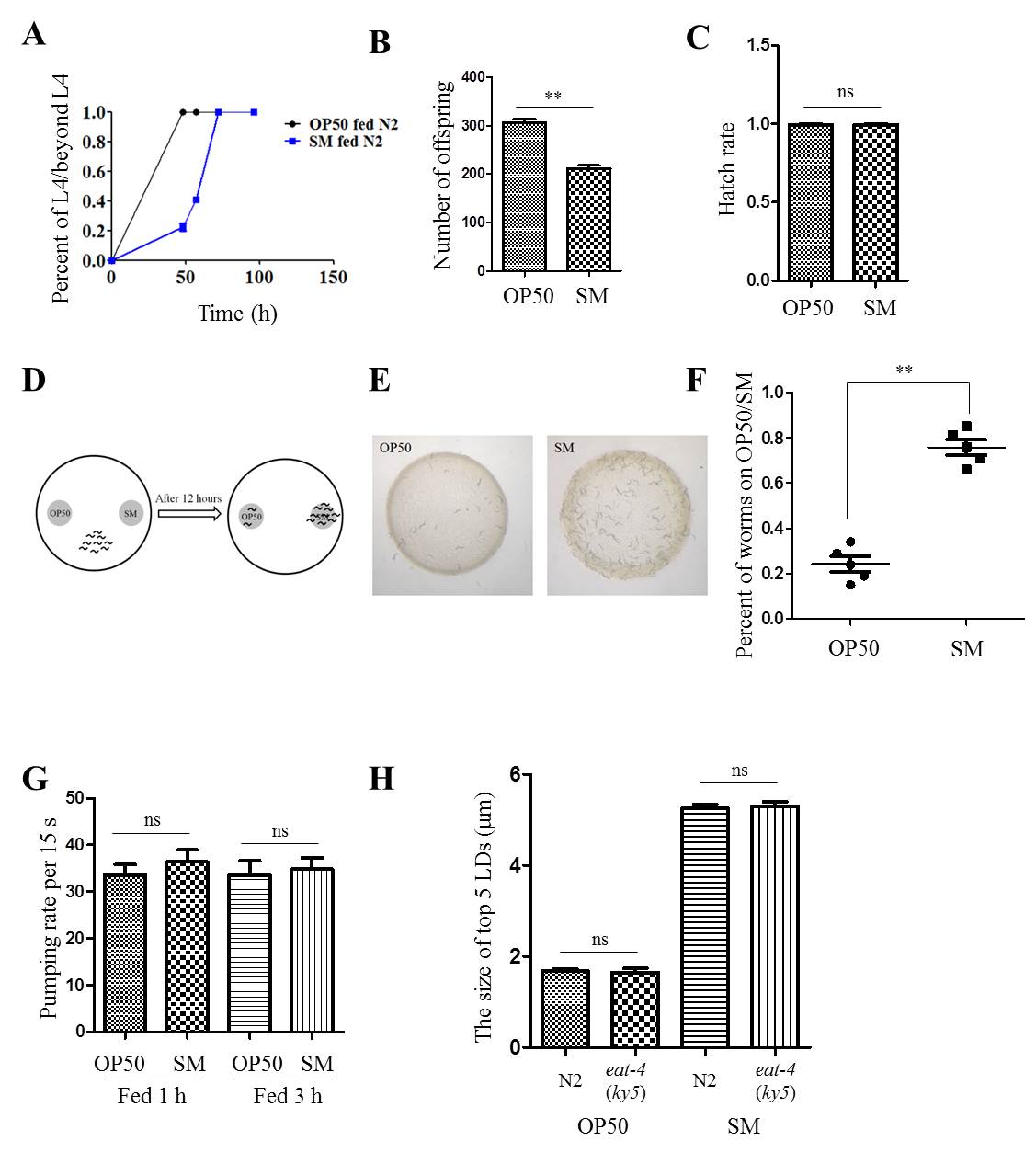

**Figure S3 Physiological measurements in *C. elegans* fed OP50 or *Stenotrophomonas maltophilia.***

**(A)** Statistics of growth rate in OP50 and *S. maltophilia*-fed N2. **(B)** Statistics of number of offspring in OP50 and *S. maltophilia*-fed N2. (n=5 for each independent experiment, ***P*<0.01, student *t*-test). **(C)**. Statistics of hatch rate in OP50 and *S. maltophilia*-fed N2. (n=5 for each independent experiment, ns, no significance, student *t*-test). **(D)** Schematic representation of the assay for the selectivity of nematodes for different diets. **(E)** The results of the selectivity of worms for OP50 or *S. maltophilia.* **(F)** Quantification of the selectivity of nematodes for OP50 or *S. maltophilia.* (n=5 for each independent experiment, ***P*<0.01, student *t*-test). **(G)** Quantification of the pumping rate of N2 worms after OP50 or *S. maltophilia* feeding for 1 h and 3 h. (n=5 for each independent experiment, ns, no significance, student *t*-test). **(H)** Quantification of the diameter of LDs. (n=5 for each independent experiment, ns, no significance, student *t*-test).

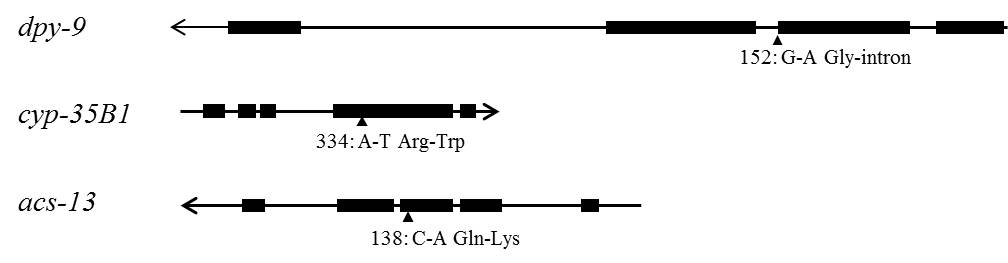

**Figure S4 Schematic representation of the gene structures and mutation sites of *dpy-9*, *cyp-35B1*, and *acs-13*.**

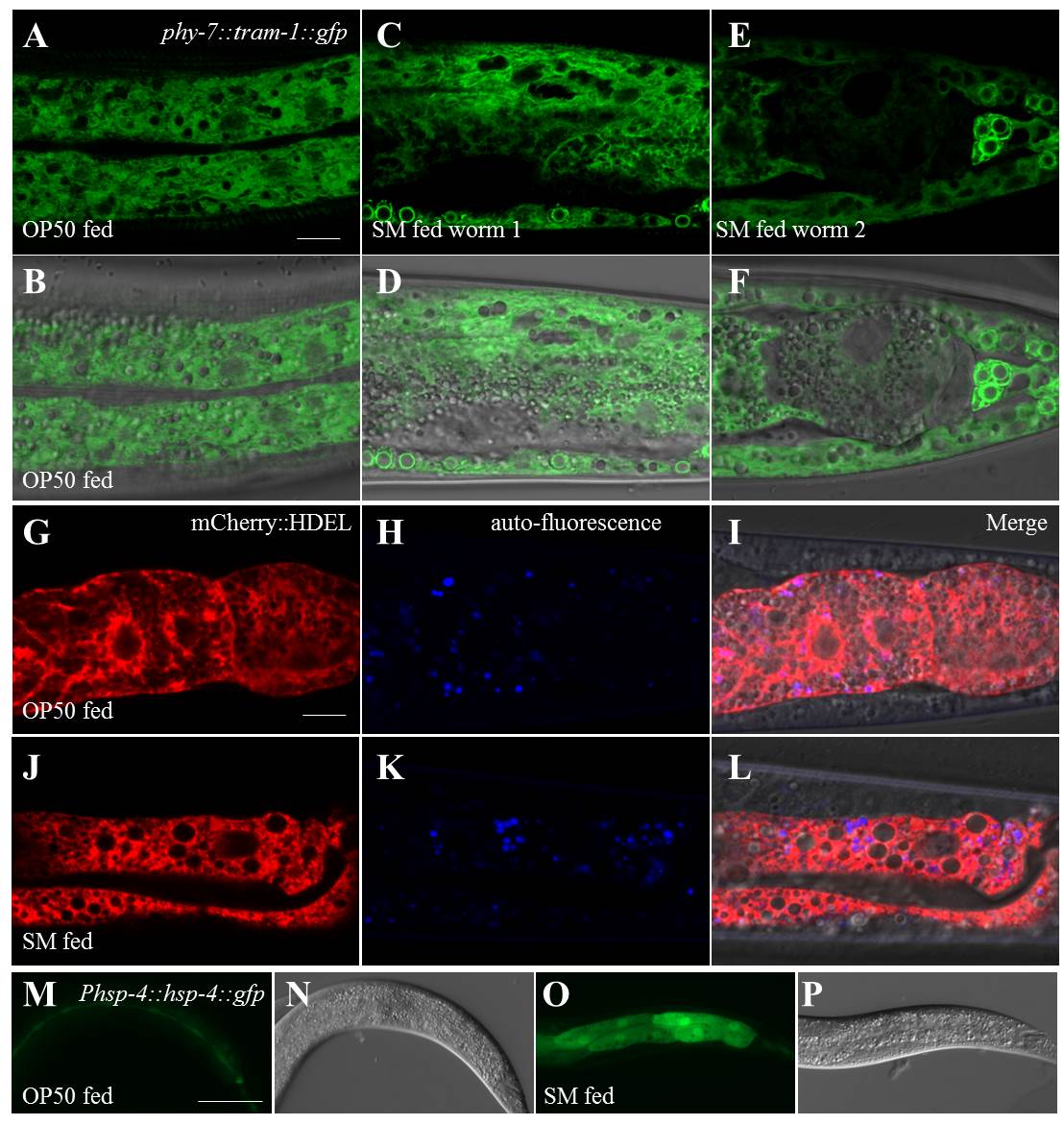

**Figure S5 *Stenotrophomonas maltophilia* induces the ER association with LDs and ER stress.**

**(A)** Fluorescence micrographs of *Phyp-7::tram-1::gfp* in OP50 fed worms. Scale Bar, 5 μm. **(B)** As in (A), but merged with DIC. **(C)** Fluorescence micrographs of *Phyp-7::tram-1::gfp* in *S. maltophilia-*fed worms. **(D)** As in (C), but merged with DIC. **(E)** As in (C), the repeated experiment results. **(F)** As in (D), the repeated experiment results. **(G)** Fluorescence micrographs of mCherry::HDEL in OP50 fed worms. Scale Bar, 5 μm. **(H)** Fluorescence micrographs of auto-fluorescence in OP50 fed worms. **(I)** As in (G), but with (H) and DIC merged. **(J)** Fluorescence micrographs of mCherry::HDEL in *S. maltophilia*-fed worms. **(K)** Fluorescence micrographs of auto-fluorescence in *S. maltophilia*-fed worms. **(L)** As in (J), but with (K) and DIC merged. **(M)** Fluorescence micrographs of *Phsp-4::hsp-4::gfp* in OP50-fed worms. Scale Bar, 5 μm. **(N)** DIC images of LDs in OP50-fed worms. **(O)** Fluorescence micrographs of *Phsp-4::hsp-4::gfp* in *S. maltophilia*-fed worms. **(P)** DIC images of LDs in *S. maltophilia*-fed worms.

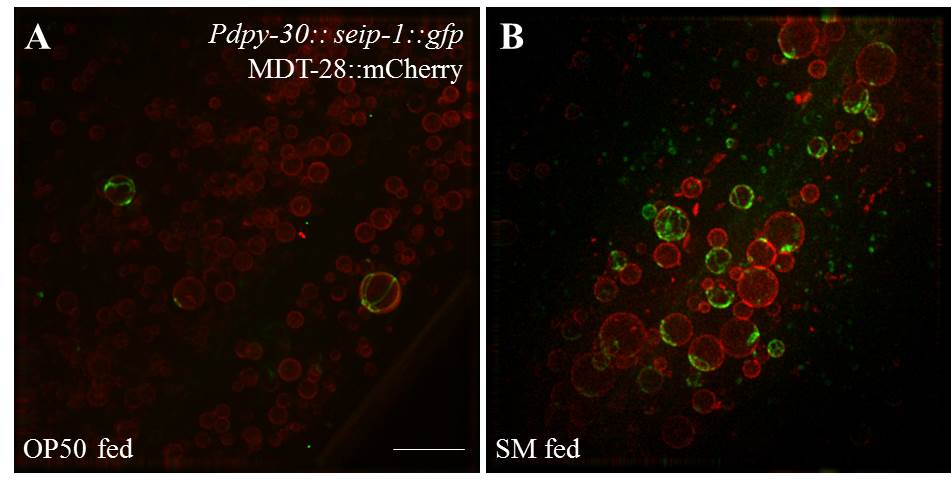

**Figure S****6 *Stenotrophomonas maltophilia* increases the percent of SEIP-1::GFP-labeled ER enwrapped LDs in *hjSi189* strain worms.**

**(A)** SIM 3D images of SEIP-1::GFP and MDT-28::mCherry in OP50 fed *hjSi189* strain worms. Scale Bar, 5 μm. **(B)** As in (A), but fed with *S. maltophilia.*
